# Localized measurements of water potential reveal large loss of conductance in living tissues of maize leaves

**DOI:** 10.1101/2023.06.06.543905

**Authors:** Piyush Jain, Annika E. Huber, Fulton E. Rockwell, Sabyasachi Sen, N. Michele Holbrook, Abraham D. Stroock

## Abstract

The water status of the living tissue in leaves between the xylem and stomata (outside xylem zone - OXZ) play a critical role for plant function and global mass and energy balance but has remained largely inaccessible. We resolve the local water relations of OXZ tissue using a nanogel reporter of water potential (*ψ*), AquaDust, that enables an in-situ, non-destructive measurement of both *ψ* of xylem and highly localized *ψ* at the terminus of transpiration in the OXZ. Working in maize, these localized measurements reveal gradients in the OXZ that are several fold larger than those based on conventional methods, and values of *ψ* in the mesophyll apoplast well below the macroscopic turgor loss potential. We find a strong loss of hydraulic conductance in both the bundle sheath and the mesophyll with decreasing xylem potential but not with evaporative demand. Our measurements suggest an active role played by the OXZ in regulating the transpiration path and our methods provide novel means to study this phenomenon.

## Introduction

Terrestrial plants play a dominant role in mediating the exchange of water, carbon dioxide (CO_2_), and energy between the earth’s surface and the atmosphere. Transpiration from plants accounts for *∼* 64% of the water flux from the land surface into the atmosphere (Good et al., 2015) and is the dominant driver of the human use of fresh water through irrigated agriculture (FAO, 2017). In coupling the soil to an unsaturated atmosphere, plants conduct water across a massive range in water potential, *ψ*, from near zero in the soil to *∼ −*100 MPa in the atmosphere (Fig. 1A). A long-standing consensus holds that the majority of this drop in potential occurs across the stomata, in the vapor phase between the interior of the leaf and the atmosphere (Nobel, 1999). Plants are assumed to achieve this control of tissue water potential by regulating stomata to maintain the mesophyll, the living tissue in the leaves, within a few MPa of leaf xylem water potential (*ψ*^xyl^), and, as a result, we expect internal mesophyll conductance to be high to keep this critical tissue close to xylem water status (Honert, 1948). But, recent work suggests that passage of the transpiration stream across the outside-xylem zone (OXZ), through the bundle sheath and mesophyll to terminal sites of evaporation (Fig. 1A), can also mediate a large drop in potential; further, this drop can vary based on changes in the conductance of the OXZ (*K*_oxz_) with the water status of the leaf and evaporative demand (Cernusak et al., 2018; Scoffoni et al., 2017; Wong et al., 2022). Nonetheless, we lack information on the constitutive properties of the OXZ and understanding of their regulation – active and passive – as a function of environmental and biological stresses (Buckley, 2017, 2019; Rockwell and Holbrook, 2017). This knowledge gap hinders scientific progress on critical outstanding questions about drought stress physiology in agricultural (Corso et al., 2020; Wu et al., 2021) and ecophysiological (Buckley, 2015, 2019) contexts and our ability to design management strategies and cultivars that are efficient and resilient with respect to water-use.

**Fig. 1.**
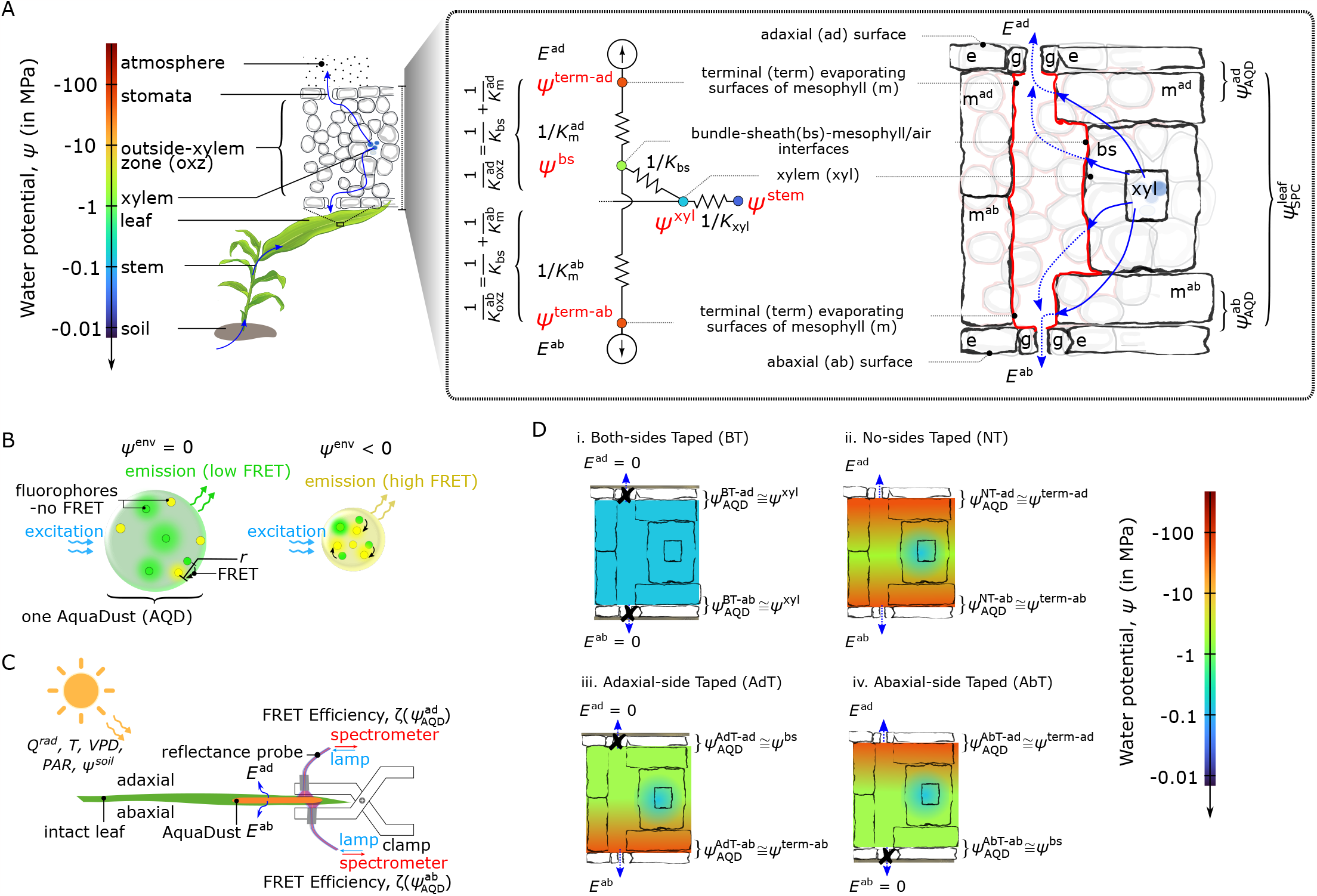
Dissection of water relations of outside xylem zone (OXZ) with AquaDust. (A) Schematic representation of soil-plant-atmosphere continuum along gradient of water potential (*ψ*) and corresponding hydraulic circuit. Expanded representation of a leaf cross-section shows the different cell types and pathways for water movement in the outside-xylem zone (OXZ). The hydraulic circuit (left) presents a simplified representation of these water paths to the adaxial (top) and abaxial (bottom) surfaces of the leaf: starting in the xylem at water potential, *ψ*^xyl^, water passes through the bundle sheath conductance (*K*_bs_) and the mesophyll conductance (*K*_m_) to reach the terminal sites of evaporation in the sub-stomatal cavity, at water potential, *ψ*^term^; water vapor then diffuses through stomata into the atmosphere with transpiration rate *E*. The nanoparticle reporter, AquaDust (Jain et al., 2021) lines the apoplastic surfaces of the mesophyll surrounding the intercellular air spaces (red lines). Its fluorescence signal reports a value of water potential from a narrow depth beneath the epidermis (*ψ*_AQD_ – as labeled), that should approximate the values of *ψ*^term^. In contrast, the Scholander Pressure Chamber provides a tissue-averaged value of water potential 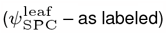. (B) Schematic diagrams showing mechanism of AquaDust nanoreporters based on Foster Resonance Energy Transfer (FRET) between pairs of dyes (green and yellow circles) in a hydrogel nanoparticle: the swollen, “wet” state when water potential in its local environment *ψ*^env^ = 0 (i.e., saturated condition), results in low FRET between donor (green circles) and acceptor (yellow circles) dye (left); and the shrunken, “dry” state when *ψ*^env^ *<* 0 (i.e., stressed condition) results in high FRET between fluorophores, thereby altering the emission spectra (right). Details on synthesis, characterization, and calibration of AquaDust is described in Jain et al. (Jain et al., 2021). (C) Schematic representation of a leaf undergoing transpiration through adaxial (*E*^ad^) and abaxial (*E*^ab^) surfaces of the leaf in a dynamic environment as a function of solar thermal radiation (*Q*^rad^) and photosynthetically active radiation (*PAR*), temperature (*T*), vapor pressure deficit (*V PD*), and stem water potential (*ψ*^stem^). Zones on the leaves infiltrated with AquaDust (orange patches on diagram) serve as reporters of the local water potential at the terminus of the OXZ in adaxial 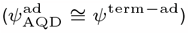 and abaxial 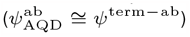 sides, captured using a short (*∼* 30 sec), minimally invasive measurement of FRET Efficiency (*ζ*(*ψ*^ab^) and (*ζ*(*ψ*^ad^) respectively) using a fiber probe positioned to collect fluorescence from the adaxial and abaxial leaf surfaces, as shown. (D) Scenarios used to manipulate the local, through-thickness gradients of water potential in leaf with selective blocking of the transpiration stream with a transparent tape: both sides obstructed (Both Tape (BT)), neither side obstructed (No Taped (NT)), and one side obstructed (Adaxial Taped (AdT) or Abaxial Taped (AbT)). These manipulations allow the water potential measured by AquaDust to provide values of water potential at different locations in the leaf by local equilibrium, as indicated with approximate equations.

Early studies using pressure probes (Frensch and Schulze, 1988; Nonami and Schulze, 1989; Shackel, 1987; Shackel and Brinckmann, 1985; Sheriff, 1982) allowed for non-invasive interrogation of OXZ water relations; however, reproducing these methods proved challenging and further use of them has not been reported. More recent studies of leaf water relations have combined whole-leaf measurements of water potential on excised tissue with the Scholander pressure chamber (SPC) and flow measurements or gas exchange to assess OXZ conductance (*K*_oxz_) (Scoffoni et al., 2017). Importantly, when applied to a transpiring leaf, the SPC provides a capacitance-weighted average over the gradient in potential in the leaf tissue (as depicted by *ψ*_SPC_ in Fig. 1A) (Cochard et al., 2013). This averaging may obscure a significant component of the water potential gradient within the OXZ, potentially leading to an overestimation of *K*_oxz_, incomplete assessments of the dependence of *K*_oxz_ on anatomy and water status (Buckley et al., 2015), and of the coupling of OXZ status to stomatal closure(Buckley and Mott, 2013), embolism avoidance (Hochberg et al., 2017), and collapse of the leaf xylem conduits (Zhang et al., 2016). Other recent studies using isolated protoplasts (Sade et al., 2014; Shatil-Cohen et al., 2011), inference from gas exchange analysis (Wong et al., 2022), and isotope discrimination (Cernusak et al., 2018; Holloway-Phillips et al., 2019) have suggested that the dynamic regulation of hydraulic conductance of the plasma membrane could decrease *K*_oxz_ to levels comparable to that of stomata. An extreme loss of the conductance along the transpiration pathway might explain observations of significant undersaturation (*<* 85% RH) at the sites of evaporation (Rockwell et al., 2022). If leaves can indeed sustain such an unsaturated state in their airspaces, it would challenge the current consensus on leaf water transport that stomata are uniquely effective at regulating water loss from leaves (Buckley, 2019; Rockwell et al., 2022).

Further progress has been hindered by the lack of practical, in-situ, non-destructive methods to resolve water potentials and conductance in the OXZ. In this paper, in leaves of maize (Zea mays L.), we use a nanogel reporter of water potential (AquaDust - Fig. 1B - (Jain et al., 2021)) in combination with gas exchange (Fig. 1C) and selective blocking of transpiration from leaf surfaces to access potentials upstream (at xylem, *ψ*^xyl^), inside (at bundle sheath-mesophyll/air interface, *ψ*^bs^), and downstream (in terminal evaporating surfaces of mesophyll adjacent to stomata, *ψ*^term^) of the OXZ (Fig. 1D). With these localized potentials, we extract values for the conductances of the leaf (*K*_leaf_), the whole OXZ (*K*_oxz_), the bundle sheath (*K*_bs_), and the abaxial mesophyll 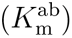 and adaxial mesophyll 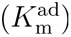 (Fig. 1B) as a function of water status (*ψ*^xyl^). These measurements and manipulations provide a new basis for the dissection of the water relations of the OXZ and open an unprecedented view of the individual contributions of the xylem, bundle sheath, and mesophyll to whole-leaf conductance and its regulation.

## Results

### Accessing localized values of water potential with AquaDust reporter

AquaDust is a nanoscale gel in which the Förster Resonance Energy Transfer (FRET) between two covalently linked dyes provides calibrated changes in the fluorescence spectra as a function of changes in water potential in the gel’s local environment (see S.I. - Sec. A and Jain et al. (Jain et al., 2021) for details). Upon infiltration through stomata into the interior of a leaf (see S.I. - Sec. B for details), AquaDust distributes itself throughout the mesophyll, coating the surfaces of the intracellular air spaces; it does not enter the symplast or the xylem (red trace in Fig. 1A; S.I. - Sec C,D and Video S1). This mode of deposition provides a first form of localization: the nanoreporters sit at the interface between the condensed phase of water in cell walls and the vapor phase that together make up the apoplast of the mesophyll.

A second form of localization comes from the optics of the fluorescent measurement: the fluorescence signal captured from the particles is localized within 15 *−* 25 *µ*m of the inner epidermal surface of the side, adaxial or abaxial, from which the measurement is performed (as labeled in Fig. 1A; Fig. 2). We interpret this localization as being due to optical absorption and scattering of both the excitation and emission light interacting with different components of the leaf tissue made up of different refractive indices. Based on these two forms of localization, we hypothesize that AquaDust measurements provide access to a local average of the water potential at the terminal sites of evaporation adjacent to stomata (*ψ*_AQD_ ≅ *ψ*^term^, Fig. 1A and Fig. 1D(ii)), the extreme end of the hydraulic path followed by transpiration from soil to the atmosphere (See S.I. - Sec. D-F for an additional explanation of methods and theory).

**Fig. 2.**
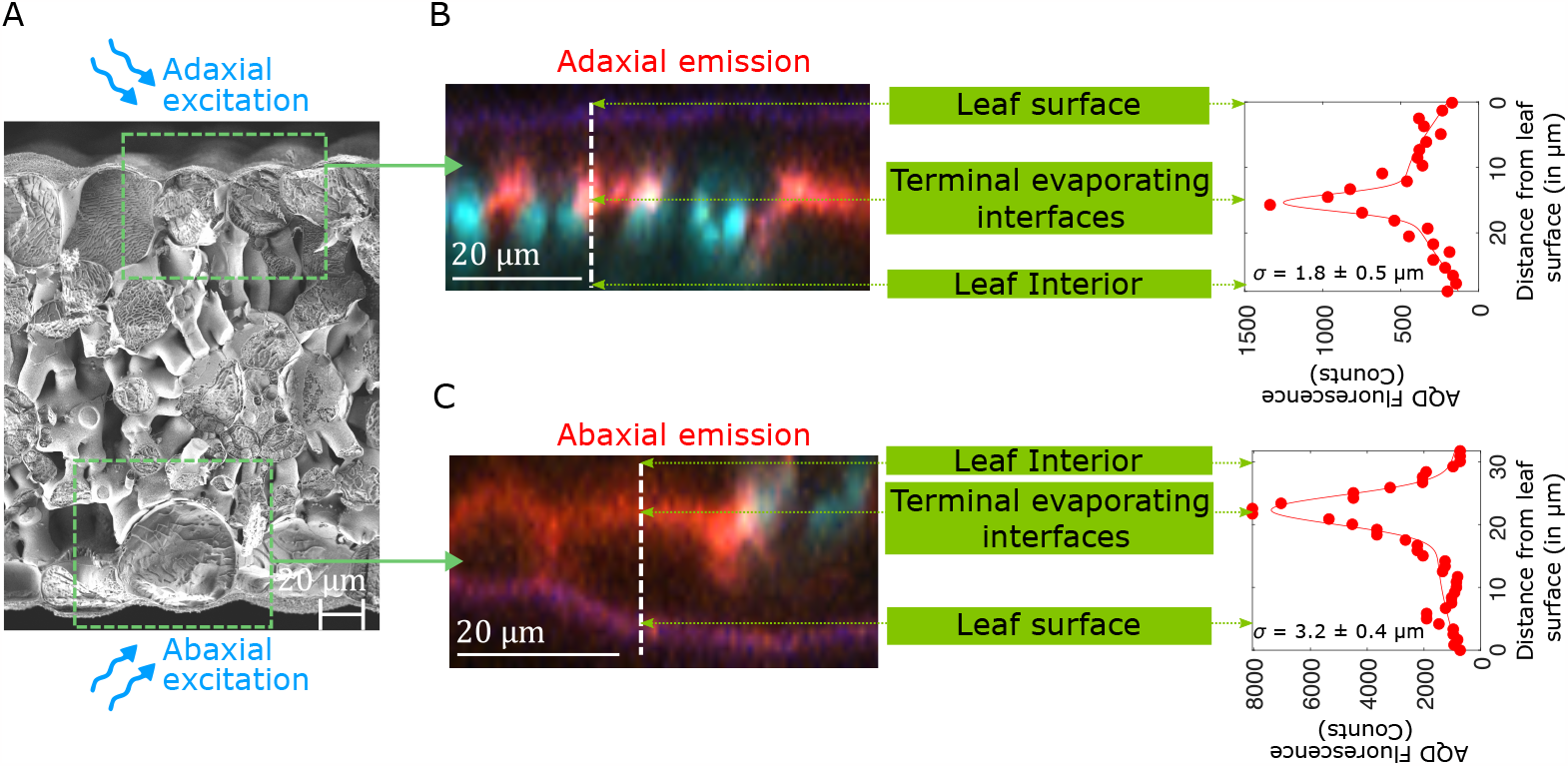
Localization of AquaDust optical signal. (A-C) Cryo-SEM micrograph of a maize leaf cross-section (A) and the depth profile of acceptor dye emission (580 *±* 10 nm) from fluorescence confocal cross-sections through adaxial (B) and abaxial (C) surfaces of maize leaf.

To access values of water potential at other locations in the cross-section of the leaf, we selectively block the transpiration from the areas of the leaf surface from which we capture the AquaDust signal (Fig. 1D). Fig. 3A presents this strategy on four segments along the length of a maize leaf: in each segment, we infiltrated AquaDust into one region on each side of the midrib; we leave four of these regions uncovered (orange regions along top half of leaf – No Tape (NT)) and cover both the adaxial and abaxial surfaces of the other four regions with a transparent tape to obstruct transpiration (shaded orange regions on the bottom half of leaf – Both Taped (BT) – See Methods or S.I. - Sec. G, Fig. S1 for details). The measurements of transpiration rate (*E*), assimilation rate (*A*), and effective stomatal conductance (*g*_s_) captured from these regions demonstrate the efficacy of the tape for suppressing gas exchange (Fig. 3B); all values dropped by *>* 90% relative to the adjacent uncovered regions. With transpiration obstructed on both sides of the leaf (BT), we assume equilibrium through the full thickness of the leaf (Fig. 1D(i)) such that 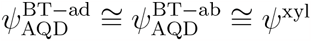, where *ψ*^xyl^ is the local potential in the xylem. In Fig. 3C, we plot the potentials reported by AquaDust in these BT regions (closed symbols); we observe agreement within uncertainty between adaxial 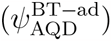 and abaxial 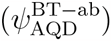 values with model predicted values of xylem water potential 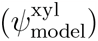 using the xylem conductance in maize based on destructive sampling (Li et al., 2009) (dashed curves – See S.I. - Sec. G and Fig. S1, details of the model are described in Jain et al. (Jain et al., 2021)). These observations provide an unprecedented basis for capturing local xylem potential, upstream of the OXZ, on intact leaves.

**Fig. 3.**
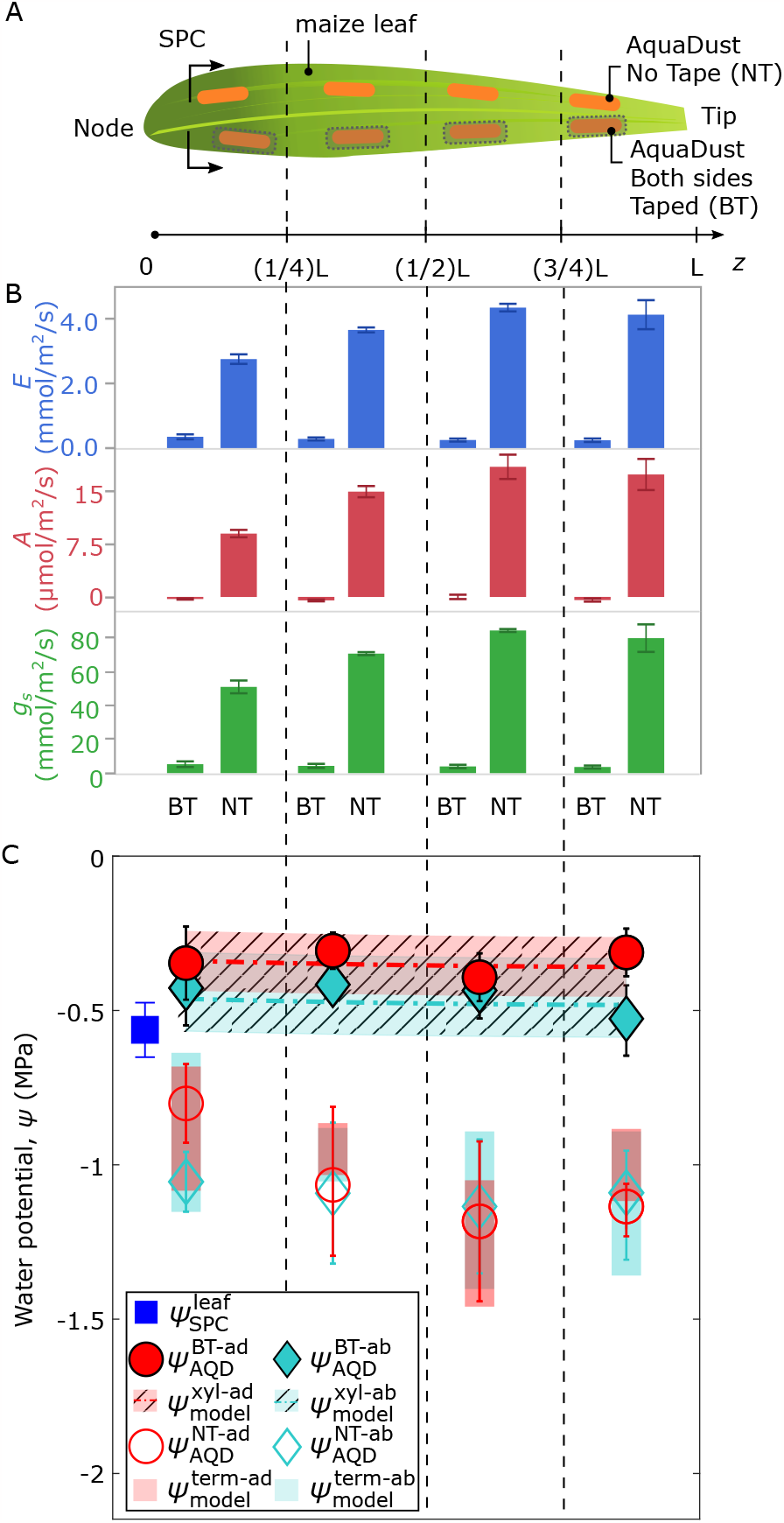
AquaDust measurements with controlled transpiration along the length of the maize leaf. (A) Diagram of maize leaf with eight regions infiltrated with AquaDust (orange) and either without any tape on either side (not-taped -NT) or covered on both sides with impermeable tape (BT). One Scholander Pressure Chamber (SPC) measurement was performed on the whole leaf (cut at indicated position) at the end of the experiment. See S.I. - Sec. G for details. (B) Transpiration rate (*E*, mmol*/*m^2^*/*s), assimilation rate (*A, µ*mol*/*m^2^*/*s), and stomatal conductance (*g*_s_, mmol*/*m^2^*/*s) in each zone on BT or NT regions (n=4, error bars represent standard error). (D) Potentials measured with AquaDust on the adaxial side for NT and BT regions (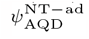 and 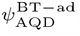) and abaxial side for NT and BT regions (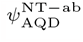 and 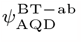) compared to model predictions of xylem water potential (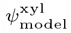- shaded) and terminal water potential (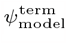- unshaded).

The open symbols in Fig. 3C present the potentials reported by AQD from the adjacent, unobstructed regions (NT) in each segment of the leaf. As indicated schematically in Fig. 1D(ii), we hypothesize that these values represent the terminal potentials at the downstream end of the OXZ during active transpiration: 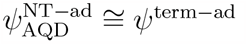 and 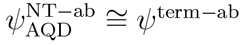. The observed values (open symbol - Fig. 3C) are consistent with this interpretation (see S.I. - Fig. S1): they fall significantly below both the values in the xylem (closed symbols – Fig. 3C) and the potential of the whole leaf captured with the SPC (blue square – Fig. 3C). Model predictions (shaded bars in Fig. 3C) based on our detailed interrogation of the variable conductance of the OXZ (see Fig. 4; S.I. - Sec. G and Table S4) agree well with these measurements of *ψ*^term^. Local access to the potentials both upstream (i.e., in BT scenarios) and downstream (i.e., in NT scenarios) of the OXZ provides a new basis for assessing the effects of transpiration on the state of water in leaf mesophyll tissue, the dominant locus of terrestrial photosynthesis.

**Fig. 4.**
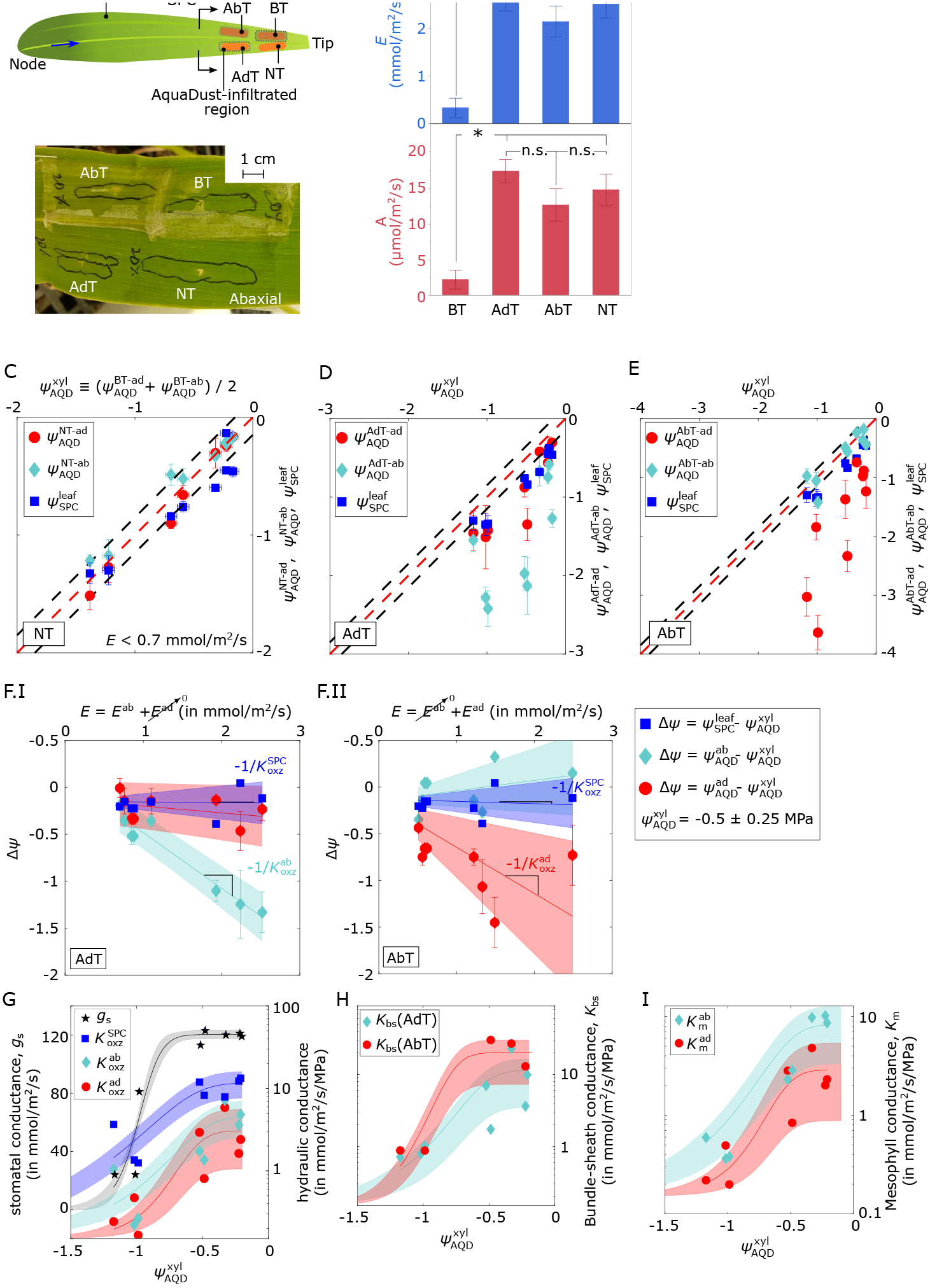
Interrogation of local leaf hydraulics with AquaDust in maize: (A) Schematic diagram and picture of a maize leaf with four different treatments to manipulate adaxial and abaxial transpiration (*E*^ad^ and *E*^ab^): suppressing transpiration (*E*) by applying tape on both sides (BT), applying tape on adaxial surface (AdT), applying tape on abaxial surface (AbT), and a control region with no tape (NT) (see Fig. 1D). Arrows indicate the approximate length of the section of leaf tissue used for leaf water potential measurement using a Scholander pressure chamber (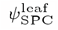). (B) Transpiration rate (*E*, mmol*/*m^2^*/*s), assimilation rate (*A, µ*mol*/*m^2^*/*s), and stomatal conductance (*g*_s_,*/* mmol m^2^ */*s) with four cases (BT, AdT, AbT, and NT) that correspond to the sites of measurements in (A) (n=4, error bars represent the standard error). See S.I. - Sec. H (Eqs. S3-S10) for the calculation of the various hydraulic conductances from these potentials. Conductances and architecture in each treatment area are as in Fig. 1A. (C-E) Measured values of whole leaf potential (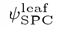– squares) and AQD potentials at adaxial (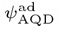 – circles) and abaxial (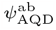 – diamonds) sides of the leaf as a function of xylem potential 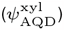 for dark-adapted leaves with low transpiration rate (*E <* 0.7 mmol*/*m^2^*/*s – C), for transpiring leaves (*E*^ab^ *>* 0.7 mmol*/*m^2^*/*s) with adaxial side taped (AdT - D), and for transpiring leaves (*E*^ad^ *>* 0.7 mmol*/*m^2^*/*s) with abaxial side taped (AbT - E). Red and dashed lines are one-to-one; black and dashed lines show uncertainty in the calibration of AquaDust (*±*0.15 MPa (Jain et al., 2021)). (F) Deviation (Δ*ψ*) of 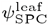 (squares), 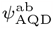 (diamonds), and 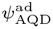 (circles) from 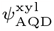 as a function of *E*^ab^ for the case of AdT zone (F.I) and *E*^ad^ for the case of AbT zone (F.II) in well-watered maize 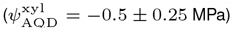. Lines represent linear regression fit, and shaded zones represent 95% confidence intervals (See S.I. - Table S2 and S3 for details). (G) Variation of stomatal conductance (*g*_s_ - black stars), effective leaf conductance (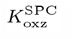 – squares – Eq. S3), adaxial and abaxial outside-xylem conductance (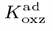 and 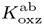 – circles and diamonds – Eqs. S5 and S6) as a function of xylem potential based on AquaDust measurement in BT region 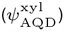. (H-I) Variation of bundle-sheath conductance *K*_bs_ (H) and of 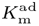 and 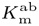, calculated for the AdT and AbT zones as a function of 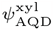 (see S.I. - Sec. H, Eqs.S3-S10, and table S4, S5, S6 for definitions of effective conductances and fits (sigmoidal curves with shaded confidence intervals)).

### Hyper-local water relations of the OXZ in maize in response to stress

In Fig. 4, we present a deeper interrogation of the water transport in the distal segment of maize leaves (Fig. 4A). When one side of a leaf is blocked while the opposite surface is transpiring, we expect that the potential of the blocked surface should be close to that of the bundle sheath surface, an expectation motivated by the prediction for distribution of *ψ* through the leaf cross-section using finite-element models of water transport in leaves (see S.I. - Sec. H, Fig. S2 and S3, (Rockwell et al., 2014a,b) for a detailed description of definitions, methods, and hydraulic architecture). To exploit this idea, we infiltrated four regions with AQD (Fig. 4A) and subjected them to four different taping treatments: (i) blocked transpiration at both surfaces (BT); (ii) unobstructed transpiration (NT); (iii) blocked transpiration at the adaxial surface only (AdT); and (iv) blocked transpiration at the abaxial surface only (AbT). We followed the response of these regions to increasing xylem stress (induced by soil drying), from near saturation to the turgor loss point (*−*1.5 *< Ψ*^xyl^ *< −*0.2 MPa, Figs. 4C-E).

As above, we interpret the doubly taped potentials as indicative of the state of the local xylem in the BT region 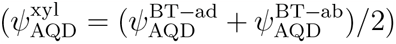. At minimal levels of transpiration (in dark, *E <* 0.7 mmol*/*m^2^*/*s), we see the convergence of the values in the NT region: 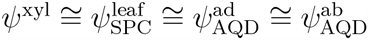 (Fig. 4C; black dashed lines represent the uncertainty of AQD measurements (*±*0.15 MPa (Jain et al., 2021)). This convergence is consistent with our expectation of near equilibrium of all components of the leaf at low transpiration fluxes, and thereby provides an in-planta validation of the accuracy of AquaDust relative to the SPC.

At higher rates of transpiration (*E >* 0.7 mmol*/*m^2^*/*s) with blocked adaxial (AdT - Fig. 4D) and blocked abaxial (AbT – Fig. 4E) surfaces, the water potentials at the transpiring surfaces deviate strongly from the xylem (red dashed, one-to-one line) as expected, but also from the SPC (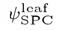 −blue squares), supporting our hypothesis that the localization of AquaDust captures a transpiration induced gradient of potential in the OXZ inaccessible to the SPC. Further support for this idea comes from leaves transpiring at varying rates at low xylem stress: the pressure gradient defined by AquaDust responds linearly to transpiration, implying a constant conductance of the OXZ in unstressed leaves, while the gradient defined by the SPC is unresponsive to transpiration (Fig. 4F.I Fig. 4F.II, see S.I. - Table S2, S3 for quantitative estimates). Finally, we observe in some cases, particularly, with declining 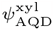 and 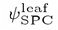, that terminal *ψ*^oxz^ is significantly more negative than the average turgor loss point for these species defined by the SPC, even though these leaves continue to have reasonable transpiration and assimilation rates, without signs of macroscopic loss of turgor (Fig. 4A).

Our localized measurements can also be used to capture the response of the OXZ to xylem stress. Using the water potential data reported in Figs. 4D and E for maize leaves at increasing levels of stress down to turgor loss, we provide the first direct assessments of a decline in the conductance of the OXZ paths to the abaxial (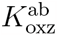, cyan diamonds) and adaxial (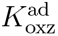, red circles) surfaces (Fig. 4G). We extract the following insights from these measurements: 1. we note that substantial symmetry between the OXZ conductances to the adaxial and abaxial surfaces, with overlapping prediction intervals (shaded cyan and pink zones) for the best fit sigmoidal responses (solid red and cyan curves) and similar values of potential at 50% loss of conductance (*ψ*_50%_ ≅ *−*0.6 MPa, see S.I. - Table S4); this symmetry is consistent with that of the morphology of the isobilateral leaves of maize (Fig. 2A); 2. we confirm the failure of conventional measurements based on the SPC (blue squares - Fig. 4G) to accurately represent the 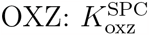 is 2 to 10-fold above that assessed with AquaDust and shows only a 3-fold drop over the range of stress studied compared to the 10-fold drop in conductance assessed using AquaDust; 3. we see in the overlaid trend in *g*_s_ (black stars) that the OXZ conductance drops earlier (at higher potentials) and further than the stomatal conductance (see S.I. - Table S4 for quantitative values of fit parameters and Fig. S4 for plots of resistances); 4. finally, we note that we can use the best-fit curves for 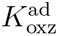 and 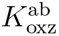 to model the variations in potential observed along the length of whole leaves (Fig. 3C - unshaded translucent red and cyan; see S.I. - Sec. Fig. S1).

The weak but resolvable deviations of potentials to the blocked surfaces opposite transpiring surfaces support our hypothesis that AquaDust signals from a non- or minimally-transpiring side of a leaf can provide access to the water potential at the bundle sheath to mesophyll transition (Fig. 1A, 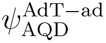- red circles in Fig. 4D; 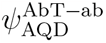- cyan diamonds in 4E; see S.I. - Sec. H, Fig. S2 for details). No existing technique provides access to these contributions in situ, yet indirect techniques (e.g., permeability measurements on protoplasts) suggest distinct regulation of water conductance by the bundle sheath and mesophyll (Shatil-Cohen et al., 2011) and molecular mechanisms for this regulation (Grunwald et al., 2022). This technique allows us to construct separate variability curves for the conductances of the bundle sheath (*K*_bs_) and mesophyll (*K*_m_) as shown in Fig. 4H-I (see S.I. - Sec. H for the definition of conductances, Table S5, S6 for values of fit parameters, and Fig. S4 B-C for plots of resistances). We note here that we only include the data from Fig. 4D-E in Fig. 4H where the deviation from 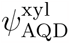 are resolvable (*> ±*0.15 MPa) using AquaDust. With this deeper dissection of the water conductance in the OXZ, we find that the both the bundle sheath and mesophyll tissues contribute to the loss of conductance with similar values of *ψ*_50%_ *∼*= 0.6 MPa, suggesting that these tissues may share both the mechanism of regulation and the functional roles by lowering of conductance in response to water stress.

## Discussion

### Interpretation and implications of terminal potentials in mesophyll and variation in OXZ conductance

AquaDust provides measurements of water potentials at the boundaries of the OXZ: the source end is defined by xylem water potential (*ψ*^xyl^), and the sink end is defined by the water potential of the terminal evaporating surfaces (*ψ*^term^) underneath the epidermis. We investigate the changes in the tissue conductance (*K*_oxz_) that couple the source and sink potentials with independent control of the environmentally relevant variables, i.e., soil water supply that defines water availability to the plant and vapor pressure deficit that defines the evaporative demand (*E*) from the plant. Access to the *ψ*^xyl^, *ψ*^term^ and *E* with varying water availability and evaporative demand help resolve important and persistent uncertainties in our understanding of the water relations of the OXZ relative to hydraulic architecture, constitutive properties, and the sites of dominant and variable conductances (Buckley, 2015; Buckley et al., 2015; Rockwell and Holbrook, 2017; Scoffoni et al., 2017).

We found that the passage of water through the OXZ creates a significant drop in *ψ*; Δ*ψ* across OXZ is of the same order or greater than that across all upstream components of the soil-plant-atmosphere continuum. Using AquaDust to access the potential at the sites of evaporation, we find that the conductance of the OXZ is *∼* 200% lower than previously measured using other techniques, with low values of *K*_oxz_ being *∼* 0.2 *−* 0.3 mmol*/*m^2^*/*MPa*/*s as measured here compared to *∼* 3 *−* 5 mmol*/*m^2^*/*MPa*/*s measured using pressure-probe method (Shackel, 1987; Shackel and Brinckmann, 1985) or *∼* 1 mmol*/*m^2^*/*MPa*/*s (Scoffoni et al., 2017; Trifiló et al., 2016) measured using SPC in cases when *ψ*^xyl^ *> Ψ*^TLP^. Notably, we observe in some cases, particularly with declining *ψ*^xyl^, that *ψ*^term^ became significantly more negative than the turgor loss point of maize, even as these leaves continued to have reasonable transpiration and assimilation rates and show no signs of macroscopic loss of turgor. While surprising, this observation aligns with recent assessments of *ψ* using indirect techniques, where significant unsaturation (*ψ*^term^ *≪ Ψ*^TLP^) was reported in leaves of *Juniperus monosperma, Pinus edulis, Gossypium hirsutum* L. and sunflower *Helianthus annuus* L. (Cernusak et al., 2018; Wong et al., 2022). Since AquaDust is localized in the apoplast, the observation that *ψ*^term^ *< Ψ*^TLP^ suggests two distinct possibilities: (1) locally, cells proximate to stomata lose turgor, or, (2) there is significant resistance at the interface between the symplast and the apoplast resulting in the breakdown of local equilibrium between water in liquid and vapor phase. Either of these possibilities has wide-ranging implications in our understanding of water transport in OXZ and requires further investigation (Rockwell et al., 2022; Wong et al., 2022).

Our direct measurements of *K*_oxz_ in physiologically relevant conditions of intact, actively assimilating and transpiring, leaves allow examination of two hypothetical scenarios explored in the literature for mechanism and functional significance of the decline in *K*_oxz_ (Scoffoni et al., 2017; Scoffoni and Sack, 2017; Shatil-Cohen et al., 2011; Trifiló et al., 2016; Wong et al., 2022). In the first, the decline in *K*_oxz_ is associated with passive, physical processes resulting from the dehydration of the OXZ tissue and morphological changes such as shrinkage and drying (with capillary failure) of cell walls (Scoffoni et al., 2017; Scoffoni and Sack, 2017; Trifiló et al., 2016); we refer to this scenario as apoplastic vulnerability, in analogy to the physical loss of conductance observed in xylem at extreme stress (typically beyond the turgor loss point). In the second scenario, the decline in *K*_oxz_ is associated with regulation of the conductance to the transpiration stream within the bundle sheath and mesophyll, likely via cell biologically controlled changes in the permeability of plasma membranes; we refer to this scenario as active regulation (Scoffoni et al., 2017; Shatil-Cohen et al., 2011; Wong et al., 2022), to emphasize its potential mechanistic and functional connections to the active processes associated with stomatal regulation.

Studies of tissue-specific contributions involving measurements of membrane permeability of protoplast (Grunwald et al., 2022; Sade et al., 2014) have linked both bundle sheath and mesophyll tissue to the decline in *K*_oxz_. Our measurements confirm that both these tissues (Fig. 4H-I) contribute to the overall decline in *K*_oxz_. Yet, the mechanisms driving declines in *K*_oxz_ remain controversial. A leading hypothesis proposes that, due to the presence of an apoplastic barrier, bundle-sheath conductance depends upon biologically regulated plasma membranes, while mesophyll conductance is dominated by an apoplastic path that contributes to declines in *K*_oxz_ by passive dehydration (Buckley et al., 2017; Scoffoni and Sack, 2017; Taneda et al., 2016). Against this latter idea, we note that the loss of hydraulic connection due to capillary failure within the cell walls at the potentials observed in *ψ*^term^ would also require a large pore size (*∼* 100 nm), an order of magnitude larger than the pore size typically observed in cell wall matrix (Dietz and Herth, 2011). Furthermore, we observe that the OXZ conductance remains relatively constant as a function of *E* (constant slope - Figs. 4F), implying that *K*_oxz_ is only a weak function of evaporative demand. This observation is inconsistent with the hypothesis that the decline in *K*_oxz_ results from the passive loss of conductance in the cell wall since this process should be more sensitive to increasing evaporative demand than xylem water potential.

Rather than being highly sensitive to VPD, we find *K*_oxz_ to be a strong function of *ψ*^xyl^ (Fig. 4G). This decline in *K*_oxz_ precedes both stomatal closure and turgor loss, favoring the hypothesis that the decrease in OXZ conductance is actively regulated within the photosynthetically active OXZ tissue to protect xylem from large tensions while keeping the stomata open for carbon uptake (Wong et al., 2022). We note that the concomitant decline in bundle-sheath and mesophyll conductance with similar values of *ψ*_50%_ (≅ 0.6 MPa) suggests a shared regulatory mechanism for water transport in both bundle-sheath and mesophyll cells. Finally, we observed that the total conductance of the OXZ falls substantially further than that of the stomata and at higher values of xylem water potential. Taken together, our measurements suggest that this active regulation of the OXZ may be an adaptive response that protects upstream components of the transpiration path and plays a role in modulating gas exchange.

## Conclusion

The local interrogation of OXZ achieved with AquaDust reveals large gradients of water potential in the outside-xylem tissue and dramatic loss of conductance of both bundle-sheath and mesophyll tissue in response to drought stress in the xylem. Our observations are consistent with the active regulation of OXZ conductance by cellular responses triggered by the declining water availability encoded in the water potential in the xylem. Our observation of apoplastic potentials well below the turgor loss point brings into question the long-standing assumption of local equilibrium between apoplast and symplast of the mesophyll tissue and the related assumption that the tissue in the OXZ may be safely represented as homogeneous continuum tissue-level properties. The large decline in the conductance of OXZ has implications for coupled processes of water loss and CO_2_ assimilation that are hosted by OZX, and point to a possible basis for non-stomatal regulation of gas exchange. This study presents a new set of tools for further interrogation of these OXZ phenomena in the contexts of crop improvement, ecophysiology, and land-atmosphere modeling in a changing climate.

## Methods

AquaDust nanoparticles (30-100 nm (range) diameter) are formed of hydrogel containing two distinct fluorescence dyes covalently linked to the matrix (Fig. 1B. See S.I. - Sec. A for details) (Jain et al., 2021). The particles swell and shrink with changes in the water potential of their local environment such that their fluorescence spectrum changes in a reproducible, calibrated manner based on Förster Resonance Energy Transfer (FRET) between the dye species.

To allow calculation of OXZ conductances (*K*_oxz_, *K*_m_, and *K*_bs_) based on Ohm’s law and hydraulic circuits, as shown in Figs. 1A, we need localized measurements of *ψ* at multiple locations along the transpiration path from the xylem to the substomatal cavities. Upon infiltration into the leaf mesophyll, AquaDust becomes distributed throughout the mesophyll on surfaces of the intracellular air spaces, as shown schematically with the red outline in Fig. 1B (see S.I. - Sec. B and Supplementary video - S1). While the particles are distributed throughout the thickness of the leaf, the intensity of fluorescence emissions collected from AquaDust peaks just beneath the epidermal layer, with the emission peak localized between *∼* 15 *−* 25 *µ*m beneath the leaf surface (as labeled in Fig. 1B and experimental demonstration in Fig. 2). We interpret this localization as being due to optical absorption and scattering of both the excitation and emission light interacting with different components of the leaf tissue made up of different refractive indices. Based on this localization, we interpret the water potentials reported by AquaDust on a transpiring leaf surface to be that of the terminal evaporating surfaces adjacent to the substomatal cavity, *ψ*^term^ (Fig. 2). AquaDust thus reports the terminus of the liquid path of transpiration, the downstream boundary condition of the OXZ.

Access to *K*_oxz_ further requires a measurement of the water potential of local xylem (*ψ*^xyl^) on the upstream side of the OXZ. We hypothesize that as the local transpiration flux approaches zero (where local refers to the surface area over which our optical probe collects AquaDust signal), the local gradient in water potential through the OXZ vanishes such that the water potential reported by AquaDust approaches the local value in the xylem: 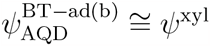 (Rockwell et al., 2014b). To access local xylem potential, we thus perform measurements of AquaDust on regions of the leaf in which we have blocked transpiration at both the adaxial and abaxial surfaces with transparent, vapor-impermeable plastic films. In maize, support comes from the convergence under low transpiration of AquaDust from both blocked and unblocked leaf surfaces with whole leaf pressure chamber measurements of water potential (see S.I. - Sec. F,G, H for details). Further validation of this hypothesis is shown in figures Fig. 3A-C and Fig. 4A-E.

## Acknowledgments and Funding

We thank Jacob L. Wszolek (Cornell Guterman Laboratory) for maintaining the plants in the greenhouse and growth chamber. This work was supported by the US Department of Agriculture National Institute of Food and Agriculture - Agriculture and Food Research Initiative Competitive Grant 2017-67007-25950; and Air Force Office of Scientific Research Grant FA9550-18-1-0345. This research is also performed in part at the Center for Research on Programmable Plant Systems and is supported by the National Science Foundation under Grant No. DBI-2019674. Further funding was provided by the Harvard MRSEC, DMR-2011754, and a Star-Friedman Challenge award (Harvard University). Imaging data were acquired through the Cornell Institute of Biotechnology’s Imaging Facility, with NIH S10OD018516 funding for the shared Zeiss LSM880 confocal/multiphoton microscope. Cryo-imaging was performed in part at the Harvard University Center for Nanoscale Systems (CNS); a member of the NNCI supported by the National Science Foundation under NSF award no. ECCS-2025158. This work was also performed in part at the Cornell NanoScale Facility, an NNCI member supported by NSF Grant NNCI-2025233.

## Author Contributions

P.J., A.E.K., F.E.R., N.M.H., and A.D.S. designed research; P.J., A.E.K., and S.S. performed research; P.J., A.E.K., F.E.R., N.M.H. and A.D.S analyzed data; and P.J., A.E.K., F.E.R., N.M.H. and A.D.S. wrote the paper.

## Declaration

Co-authors P.J. and A.D.S. are listed as inventors on Patent Number: US11536660B2 titled ‘In situ sensing of water potential’ filed by Cornell University.

